# Recombinase Polymerase Amplification Assay for the Field Detection of Mal Secco Disease by *Plenodomus tracheiphilus*

**DOI:** 10.1101/2023.10.21.563392

**Authors:** Ermes Ivan Rovetto, Matteo Garbelotto, Salvatore Moricca, Marcos Amato, Federico La Spada, Santa Olga Cacciola

**Affiliations:** Department of Agriculture, Food and Environment, University of Catania, 95123 Catania, Italy; Department of Environmental Science, Policy and Management, University of California, Berkeley, CA 94720, United States; Department of Agri-food Production and Environmental Sciences, Plant Pathology and Entomology Division - University of Florence, Piazzale delle Cascine 28, 50144 – Firenze, Italy; Agdia EMEA, 31 rue de Seine 91450 Soisy-sur-Seine, France

**Keywords:** Recombinase polymerase amplification, molecular detection, Real-Time PCR, mal secco, vascular disease, *Plenodomus tracheiphilus*, *Citrus limon*, mitosporic fungi, quarantine pathogen, EPPO list

## Abstract

In this study, we developed a new diagnostic assay based on the recombinase polymerase amplification (RPA) technology to detect *Plenodomus tracheiphilus*, the anamorphic fungus responsible for the destructive vascular disease of lemon named mal secco, in infected tissues of host plants. A 142 bp RPA-compatible barcode was sought within the 544 bp Internal Transcriber Spacer (ITS) fragment identified in a previous study and its *P. tracheiphilus*-specificity was confirmed by BLAST in the NCBI database. This was the premise to design an RPA probe (RPA_Ptrach_Probe). The specificity and inclusivity of the RPA assay were tested on gDNA isolated from tissues of *C. limon*, isolates of *P. tracheiphilus* of various origins and axenic cultures of non-target organisms, including fungal and oomycete pathogens typically associated to citrus trees, such as *Alternaria* spp., *Colletotrichum* spp., *Phyllosticta* spp*., Penicillium* spp., *Phytophthora* spp. With a detection threshold of 1.0 pg of gDNA the RPA assay proved to be as sensitive as the SYBR® Green I Real Time-PCR test included in the diagnostic protocol for *P. tracheiphilus* of the European and Mediterranean Plant Protection Organization. RPA assay was even more sensitive than Real Time-PCR in tests on DNA samples obtained through a rapid extraction method. In tests, on naturally infected lemon twigs, molecular approaches were comparable to each other and performed better than conventional isolation method. Overall, results of this study demonstrate the potential of RPA for rapid, easy to handle and cost effective in-field diagnosis of mal secco.

## 1. Introduction

Mal secco of citrus, caused by the mitosporic fungus *Plenodomus tracheiphilus* (Petri) Gruyter, Aveskamp, and Verkley (formerly *Phoma tracheiphila* (Petri) Kantschaveli and Gikashvili), in the family *Leptosphaeriaceae*, order *Pleosporales* (de Gruyter *et al*., 2013), is one of the most relevant diseases affecting citrus cultivations (Migheli *et al*., 2009). It is a devastating vascular disease, which primarily occurs in lemon (*Citrus limon* (L.) Burm. f.) and, to a lesser extent, in citron, bergamot, lime, sour orange, and rough lemon (Licciardello *et al*., 2006; Migheli *et al*., 2009; Nigro *et al*., 2015). Mal secco of lemon is present throughout most of the Mediterranean region, including the Black Sea area, with the exception of a few countries (Abbate *et al*., 2019; Demontis *et al*., 2008).

The European and Mediterranean Plant Protection Organization (EPPO) has classified *P. tracheiphilus* as an A2 quarantine pest. The pathogen is also considered a quarantine concern by several regional plant protection agencies worldwide, including the Asia and Pacific Plant Protection Commission (APPPC), the Caribbean Plant Protection Commission (CPPC), the Comité Regional de Sanidad Vegetal para el Cono Sur (COSAVE), and the North American Plant Protection Organization (NAPPO) (Licciardello *et al*., 2006; Migheli *et al*., 2009; Nigro *et al*., 2011).

The most effective strategies for the management of mal secco include several actions, such as the adoption of preventive measures, the planning of phytosanitary programs, and the early diagnosis of the disease (El boumlasy *et al*., 2022; Demontis *et al*., 2008; Licciardello *et al*., 2006). According to the EPPO diagnostic protocol number PM 7/048 for (EPPO, 2015), the diagnosis of malsecco disease is considered positive when the pathogen is identified in potentially infected plants either by direct isolation or by molecular detection from plant tissues. Currently, the molecular methods approved by EPPO for the diagnosis of mal secco are a polymerase chain reaction (PCR)-based *P. tracheiphilus-*specific assay (Balmas *et al*., 2005) and a quantitative TaqMan® / SYBR® Green I real-time PCR protocol (Demontis *et al*., 2008). In spite of the well-known advantages of PCR-based technologies, such as their reliability resulting from years of application, a formidable sensitivity, high specificity, and increased sample throughput (Demontis *et al*., 2008; Ray *et al*., 2017; Wong and Medrano, 2005), such technologies have limitations in terms of cost, complexity and can be unsuitable for disease diagnosis in the field (Wong and Medrano, 2005). In the last decade, several new molecular methods have been developed: isothermal techniques are particularly noteworthy, given they do not require thermal cycling equipment for the amplification of nucleic acids (Crannell *et al*., 2014). Within the options of isothermal technologies developed for the detection of pathogens, the recombinase polymerase amplification (RPA) stands out for its simplicity, specificity, high sensitivity, and rapidity (Lobato and O’Sullivan, 2018; Piepenburg *et al*., 2006). This technology is based on a recombinase-dependent hybridization of an oligonucleotide primer pair with a double-stranded (ds) DNA target fragment (Ivanov *et al*., 2021). Three core enzymes are employed in RPA: i) a recombinase, ii) a single-stranded DNA-binding protein (SSB), and iii) a strand-displacing polymerase. Specifically, in the amplification process, the recombinase binds itself to primers thus generating a complex recombinase/primers chimera that is then paired with homologous sequences present in the dsDNA target fragment. Then, the SSB binds to the complex formed by the recombinase/primers and prevents its displacement. Finally, the strand-displacing polymerase begins the DNA synthesis starting from the point where the primers have bound to the target fragment (Lobato and O’Sullivan, 2018; Piepenburg *et al*., 2006). The RPA exponential amplification is accomplished by the cyclic repetition of this process, which proceeds at a low and constant temperature, typically comprised in the range 25-42 °C (Ivanov *et al*., 2021; Piepenburg *et al*., 2006). The thermal requirements of RPA, can be easily satisfied using a simple thermal assay device, even portable, make the RPA both technically easy and cost-effective (Daher *et al*., 2016; Li *et al*., 2019; Lillis *et al*., 2014; Lobato and O’Sullivan, 2018). The high specificity and sensitivity of this technology is guaranteed by the strict requirements of the primers. In this respect, the optimal formation of the complex recombinase/primers is ensured by a primer length of 28–35 nucleotides each; additionally, primers GC content should be between 30% and 70% and tandem repeats of nucleotides should be avoided (Cesbron *et al*., 2023; Daher *et al*., 2016). An additional strength of RPA is represented by the chemistry related to the detection of the amplicons generated. The most performing RPA systems include the use of probes presenting a structure including a tetrahydrofuran (THF) abasic–site flanked by nucleotides modified with a fluorophore (usually the fluorine FAM) and a quencher; a block at the 3′-end is also included to prevent the probe from acting as an amplification primer. This particular chemistry of the RPA probe provides an additional specificity-proofreading to the assay. In fact, the specific pairing of the probe to the complementary DNA activates an endonuclease which cuts the probe in correspondence of the THF, leading to a measurable increase in fluorescence (Piepenburg *et al*., 2006). Other strengths of RPA include a high tolerance of impure samples and, under a practice point of view, the availability of reagents in form of lyophilized pellets. These additional features make this technique markedly suitable for field applications (Daher *et al*., 2016). In light of these advantages, RPA has been successfully used for the development of assays for the early detection of several plant pathogens, including viruses (Babu *et al*., 2018; Kapoor *et al*., 2017; Mekuria *et al*., 2014; Silva *et al*., 2015; Zhang *et al*., 2014), bacteria (Buddhachat *et al*., 2022; Cesbron *et al*., 2023; Strayer-Scherer *et al*., 2019), oomycetes (Guo *et al*., 2023; Lu *et al*., 2021; Miles *et al*., 2015; Munawar *et al*., 2019, 2020; Rojas *et al*., 2017; Yu *et al*., 2019) and fungi (Changtor *et al*., 2023; Chen *et al*., 2022; Ju *et al*., 2020; Xu *et al*., 2023; Zhao *et al*., 2021).

Due to the importance of mal secco disease, whose management needs to be supported by rapid, user-friendly, and cost-effective diagnostic tools, this study aimed to develop a *P. tracheiphilus*-RPA assay. The sensitivity of the RPA assay was evaluated in comparison to the EPPO standard SYBR^®^ Green I Real Time-PCR assay of Demontis *et al*. (2008). Finally, the suitability of the *P. tracheiphila*-RPA assay for field applications was evaluated on plant samples collected from both mal secco-symptomatic and non-symptomatic lemon plants.

## 2. Results

Experiments carried out in this study led to the development of an RPA assay for the detection of *P. tracheiphilus*. Specifically, as a first step, a 142 bp RPA technology-compatible barcode within the Internal Transcriber Spacer (ITS) of *P. tracheiphilus* was selected based on a consensus alignment of *P. tracheiphilus* ITS sequences deposited in NCBI (Figure 1). BLAST searches of the NCBI nucleotide database confirmed *in silico* that only *P. tracheiphilus* sequences had a 100% homology with the chosen barcode, while the second closest match was that of *Plenodomus collinsoniae* with a 93,33% homology.

**Figure 1.**
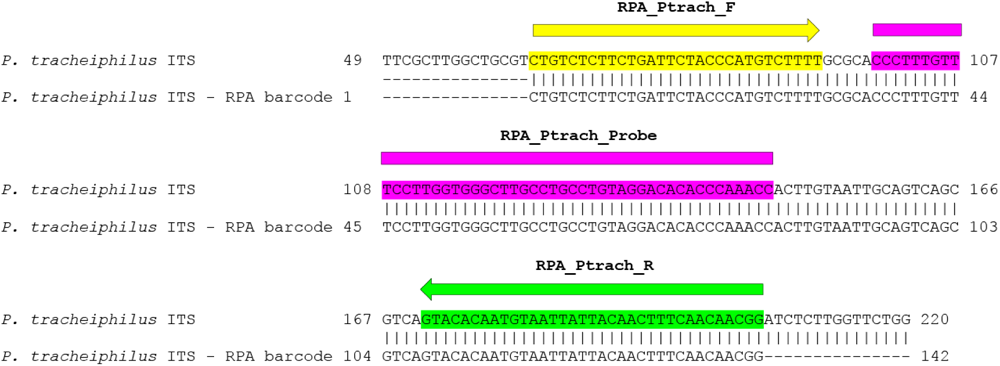
Alignment of consensus sequences of the ITS region of *Plenodomus tracheiphilus* (Genbank accession numbers: AY531665, AY531666, AY531667, AY531668, AY531669, AY531670, AY531671, AY531672, AY531673, AY531674, AY531675, AY531676, AY531677, AY531678, AY531679, AY531680, AY531681, AY531682, AY531689) and binding sites of primers and probe.

Then, a primer couple best matching RPA amplification requirements was designed (Figure 1, Table 1) and its *P. tracheiphilus*-specificity was preliminarily confirmed by both *in silico* and conventional PCR amplifications.

**Table 1.**
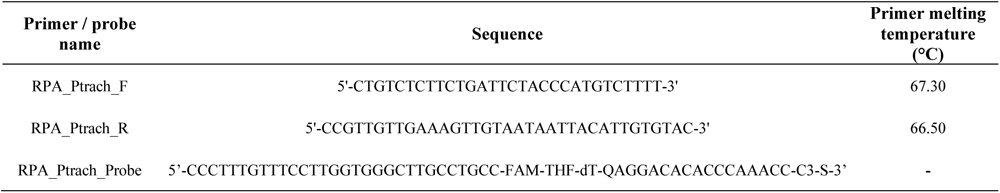
Sequences of primers and probe designed in this study for performing *Plenodomus tracheiphilus*-selective RPA amplifications.

Finally, an RPA probe was designed to complete the assay (Figure 1, Table 1).

### 2.1. Plenodomus tracheiphilus-specificity and -inclusivity of RPA assay

*Plenodomus tracheiphilus*-specificity of the developed RPA assay was tested on gDNA isolated from stem fragments of *C. limon* and pure cultures of 11 specimens of non-target organisms. Non-target organisms included fungal and oomycete pathogens typically associated to citrus cultivation.

Neither the gDNA from plant material nor non-target organisms produced RPA amplification, confirming the absence of any cross reaction with non-target DNA (Table 2). Inclusivity of the RPA assay was tested on DNA extracted from 29 specimens of the target organism (*P. tracheiphilus*). All strains of *P. tracheiphilus* were detected in RPA runs, confirming the whole inclusivity of the developed assay (Table 2).

**Table 2.**
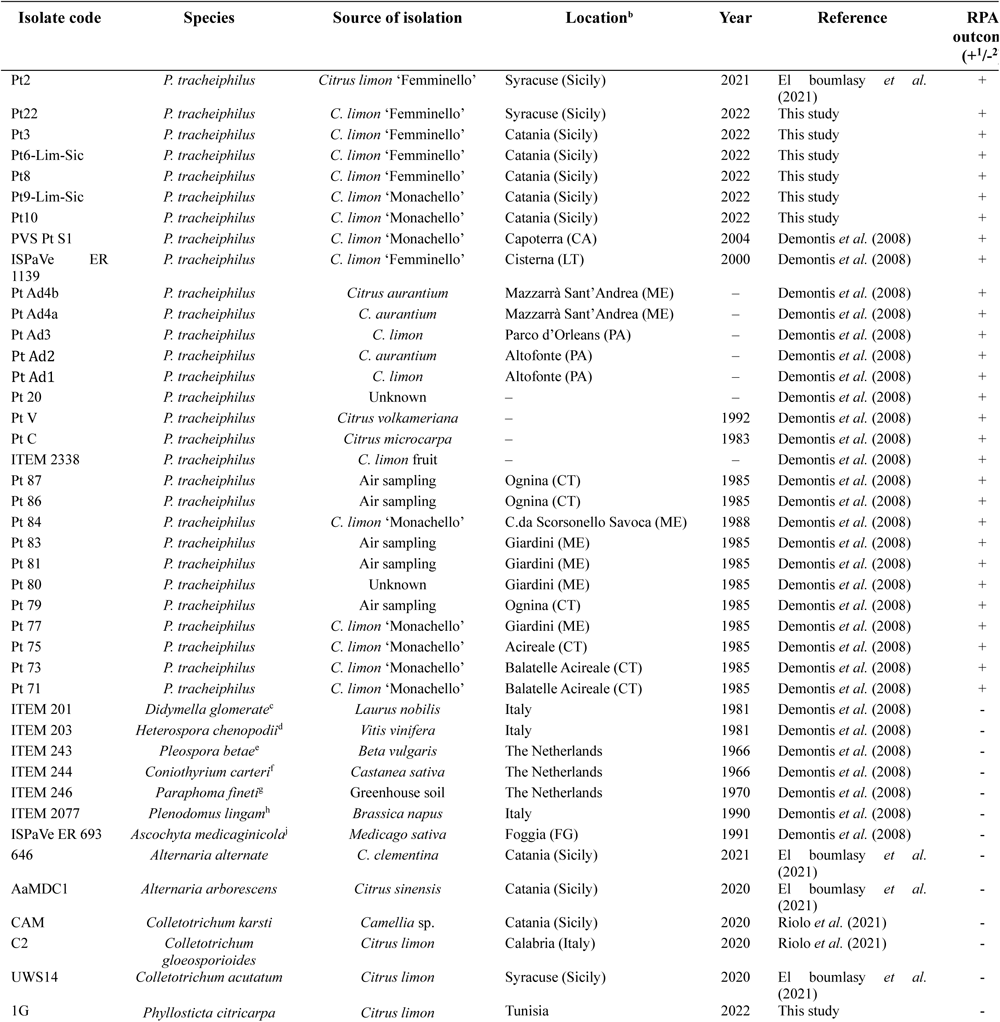
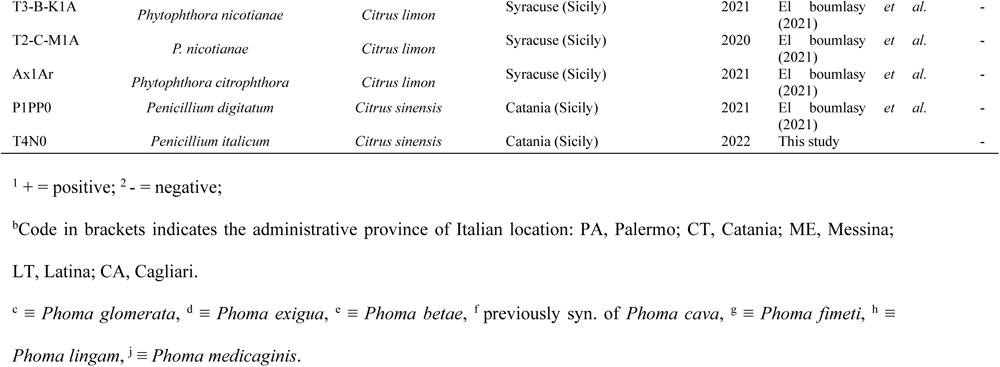
Isolate code, species, source of isolation, location, year of recovery, reference work, and RPA outcome of fungal and oomycete isolates used in this study.

| Isolate code | Species | Source of isolation | Location<sup>b</sup> | Year | Reference | RPA outcome (+<sup>1</sup>/-<sup>2</sup>) |
| --- | --- | --- | --- | --- | --- | --- |
| Pt2 | *P. tracheiphilus* | *Citrus limon* 'Femminello' | Syracuse (Sicily) | 2021 | El boumlasy *et al.* (2021) | + |
| Pt22 | *P. tracheiphilus* | *C. limon* 'Femminello' | Syracuse (Sicily) | 2022 | This study | + |
| Pt3 | *P. tracheiphilus* | *C. limon* 'Femminello' | Catania (Sicily) | 2022 | This study | + |
| Pt6-Lim-Sic | *P. tracheiphilus* | *C. limon* 'Femminello' | Catania (Sicily) | 2022 | This study | + |
| Pt8 | *P. tracheiphilus* | *C. limon* 'Femminello' | Catania (Sicily) | 2022 | This study | + |
| Pt9-Lim-Sic | *P. tracheiphilus* | *C. limon* 'Monachello' | Catania (Sicily) | 2022 | This study | + |
| Pt10 | *P. tracheiphilus* | *C. limon* 'Monachello' | Catania (Sicily) | 2022 | This study | + |
| PVS Pt S1 | *P. tracheiphilus* | *C. limon* 'Monachello' | Capoterra (CA) | 2004 | Demontis *et al.* (2008) | + |
| ISPaVe ER 1139 | *P. tracheiphilus* | *C. limon* 'Femminello' | Cisterna (LT) | 2000 | Demontis *et al.* (2008) | + |
| Pt Ad4b | *P. tracheiphilus* | *Citrus aurantium* | Mazzarrà Sant'Andrea (ME) | – | Demontis *et al.* (2008) | + |
| Pt Ad4a | *P. tracheiphilus* | *C. aurantium* | Mazzarrà Sant'Andrea (ME) | – | Demontis *et al.* (2008) | + |
| Pt Ad3 | *P. tracheiphilus* | *C. limon* | Parco d'Orleans (PA) | – | Demontis *et al.* (2008) | + |
| Pt Ad2 | *P. tracheiphilus* | *C. aurantium* | Altofonte (PA) | – | Demontis *et al.* (2008) | + |
| Pt Ad1 | *P. tracheiphilus* | *C. limon* | Altofonte (PA) | – | Demontis *et al.* (2008) | + |
| Pt 20 | *P. tracheiphilus* | Unknown | – | – | Demontis *et al.* (2008) | + |
| Pt V | *P. tracheiphilus* | *Citrus volkameriana* | – | 1992 | Demontis *et al.* (2008) | + |
| Pt C | *P. tracheiphilus* | *Citrus microcarpa* | – | 1983 | Demontis *et al.* (2008) | + |
| ITEM 2338 | *P. tracheiphilus* | *C. limon* fruit | – | – | Demontis *et al.* (2008) | + |
| Pt 87 | *P. tracheiphilus* | Air sampling | Ognina (CT) | 1985 | Demontis *et al.* (2008) | + |
| Pt 86 | *P. tracheiphilus* | Air sampling | Ognina (CT) | 1985 | Demontis *et al.* (2008) | + |
| Pt 84 | *P. tracheiphilus* | *C. limon* 'Monachello' | C.da Scorsonello Savoca (ME) | 1988 | Demontis *et al.* (2008) | + |
| Pt 83 | *P. tracheiphilus* | Air sampling | Giardini (ME) | 1985 | Demontis *et al.* (2008) | + |
| Pt 81 | *P. tracheiphilus* | Air sampling | Giardini (ME) | 1985 | Demontis *et al.* (2008) | + |
| Pt 80 | *P. tracheiphilus* | Unknown | Giardini (ME) | 1985 | Demontis *et al.* (2008) | + |
| Pt 79 | *P. tracheiphilus* | Air sampling | Ognina (CT) | 1985 | Demontis *et al.* (2008) | + |
| Pt 77 | *P. tracheiphilus* | *C. limon* 'Monachello' | Giardini (ME) | 1985 | Demontis *et al.* (2008) | + |
| Pt 75 | *P. tracheiphilus* | *C. limon* 'Monachello' | Acireale (CT) | 1985 | Demontis *et al.* (2008) | + |
| Pt 73 | *P. tracheiphilus* | *C. limon* 'Monachello' | Balatelle Acireale (CT) | 1985 | Demontis *et al.* (2008) | + |
| Pt 71 | *P. tracheiphilus* | *C. limon* 'Monachello' | Balatelle Acireale (CT) | 1985 | Demontis *et al.* (2008) | + |
| ITEM 201 | *Didymella glomerata*<sup>c</sup> | *Laurus nobilis* | Italy | 1981 | Demontis *et al.* (2008) | - |
| ITEM 203 | *Heterospora chenopodii*<sup>d</sup> | *Vitis vinifera* | Italy | 1981 | Demontis *et al.* (2008) | - |
| ITEM 243 | *Pleospora betae*<sup>e</sup> | *Beta vulgaris* | The Netherlands | 1966 | Demontis *et al.* (2008) | - |
| ITEM 244 | *Coniothyrium carteri*<sup>f</sup> | *Castanea sativa* | The Netherlands | 1966 | Demontis *et al.* (2008) | - |
| ITEM 246 | *Paraphoma fineti*<sup>g</sup> | Greenhouse soil | The Netherlands | 1970 | Demontis *et al.* (2008) | - |
| ITEM 2077 | *Plenodomus lingam*<sup>h</sup> | *Brassica napus* | Italy | 1990 | Demontis *et al.* (2008) | - |
| ISPaVe ER 693 | *Ascochyta medicaginicola*<sup>i</sup> | *Medicago sativa* | Foggia (FG) | 1991 | Demontis *et al.* (2008) | - |
| 646 | *Alternaria alternate* | *C. clementina* | Catania (Sicily) | 2021 | El boumlasy *et al.* (2021) | - |
| AaMDC1 | *Alternaria arborescens* | *Citrus sinensis* | Catania (Sicily) | 2020 | El boumlasy *et al.* (2021) | - |
| CAM | *Colletotrichum karsti* | *Camellia* sp. | Catania (Sicily) | 2020 | Riolo *et al.* (2021) | - |
| C2 | *Colletotrichum gloeosporioides* | *Citrus limon* | Calabria (Italy) | 2020 | Riolo *et al.* (2021) | - |
| UWS14 | *Colletotrichum acutatum* | *Citrus limon* | Syracuse (Sicily) | 2020 | El boumlasy *et al.* (2021) | - |
| 1G | *Phyllosticta citricarpa* | *Citrus limon* | Tunisia | 2022 | This study | - |
| T3-B-K1A | *Phytophthora nicotianae* | *Citrus limon* | Syracuse (Sicily) | 2021 | El boumlasy *et al.* (2021) | - |
| T2-C-M1A | *P. nicotianae* | *Citrus limon* | Syracuse (Sicily) | 2020 | El boumlasy *et al.* (2021) | - |
| Ax1Ar | *Phytophthora citrophthora* | *Citrus limon* | Syracuse (Sicily) | 2021 | El boumlasy *et al.* (2021) | - |
| P1PP0 | *Penicillium digitatum* | *Citrus sinensis* | Catania (Sicily) | 2021 | El boumlasy *et al.* (2021) | - |
| T4N0 | *Penicillium italicum* | *Citrus sinensis* | Catania (Sicily) | 2022 | This study | - |

<sup>1</sup> + = positive; <sup>2</sup> - = negative;
<sup>b</sup>Code in brackets indicates the administrative province of Italian location: PA, Palermo; CT, Catania; ME, Messina; LT, Latina; CA, Cagliari.
<sup>c</sup> ≡ *Phoma glomerata*, <sup>d</sup> ≡ *Phoma exigua*, <sup>e</sup> ≡ *Phoma betae*, <sup>f</sup> previously syn. of *Phoma cava*, <sup>g</sup> ≡ *Phoma fimeti*, <sup>h</sup> ≡ *Phoma lingam*, <sup>i</sup> ≡ *Phoma medicaginis*.

### 2.2. Sensitivity of the RPA assay vs. Demontis’s Real Time-PCR assay

The sensitivity of the RPA assay was evaluated on different kinds of samples from which DNA was isolated through four different protocols (Figure 2).

**Figure 2.**
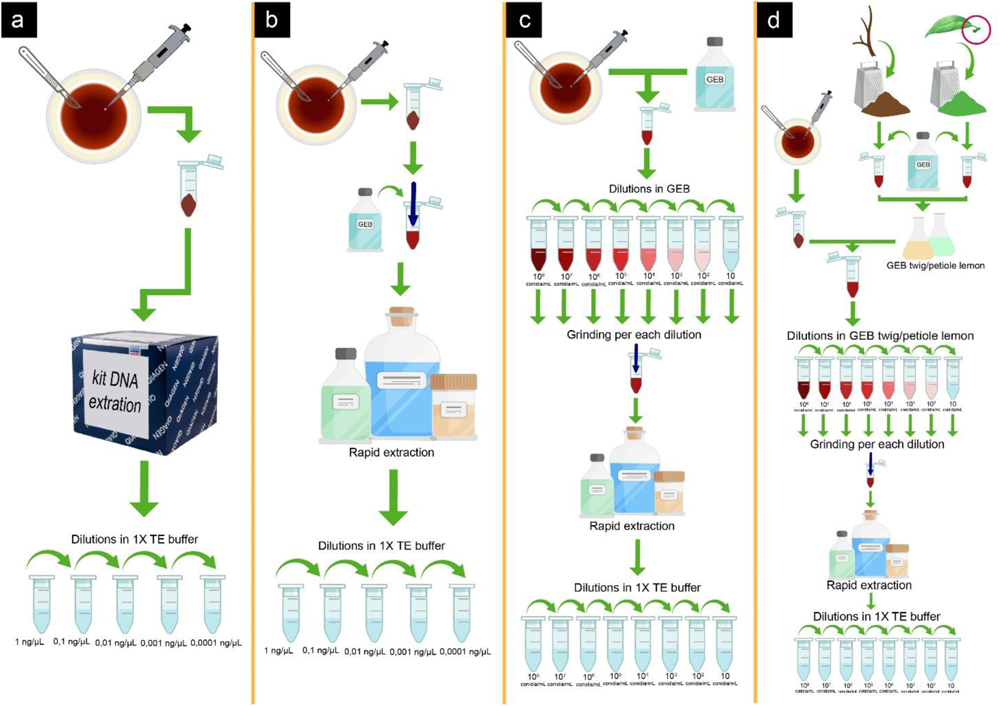
Schematic representation of the extraction and preparation process of DNA from *Plenodomus tracheiphilus* samples for the sensitivity test. In **a**. 10-fold-dilutions of purified gDNA of the *P. tracheiphilus* isolate Pt2 extracted from a bulk of 10^9^ conidia using the PowerPlant® Pro DNA Isolation Kit. In **b**. 10-fold-dilutions of non-purified gDNA of the isolate Pt2 extracted from a bulk of 10^9^ conidia using a rapid extraction protocol (according to (Edwards *et al*., 1991). In **c**. Conidia of the isolate Pt2 suspended in GEB lysis buffer (concentrations of suspension ranging from 10^8^ to 10 conidia/ml) were subjected to rapid DNA extraction (according to (Edwards *et al*., 1991); then, gDNA samples were subjected to sensitivity tests. In **d**. Conidia of the isolate Pt2 suspended in a plant crude extract (twigs/petioles powder macerated in GEB lysis buffer) (concentrations of suspension ranging from 10^8^ to 10 conidia/ml) were subjected to rapid DNA extraction (according to (Edwards *et al*., 1991); then, gDNA samples were subjected to sensitivity tests.

In ‘procedure i’ (Figure 2a) the sensitivity of the RPA was evaluated on four independent 10-folds dilutions of a 1.0 ng/µl concentrated (corresponding to about 2.85 × 10^3^ genome copies/µl) solution of purified gDNA extracted from 10^9^ conidia of the isolate Pt2 by using the PowerPlant® Pro DNA Isolation Kit. Both RPA and Demontis’s Real Time-PCR gave amplifications up to 0.001 ng of gDNA (*i.e.* 1.0 pg of gDNA, corresponding to ∼ 29 genome copies/reaction) (Figure 3, a and c).

**Figure 3.**
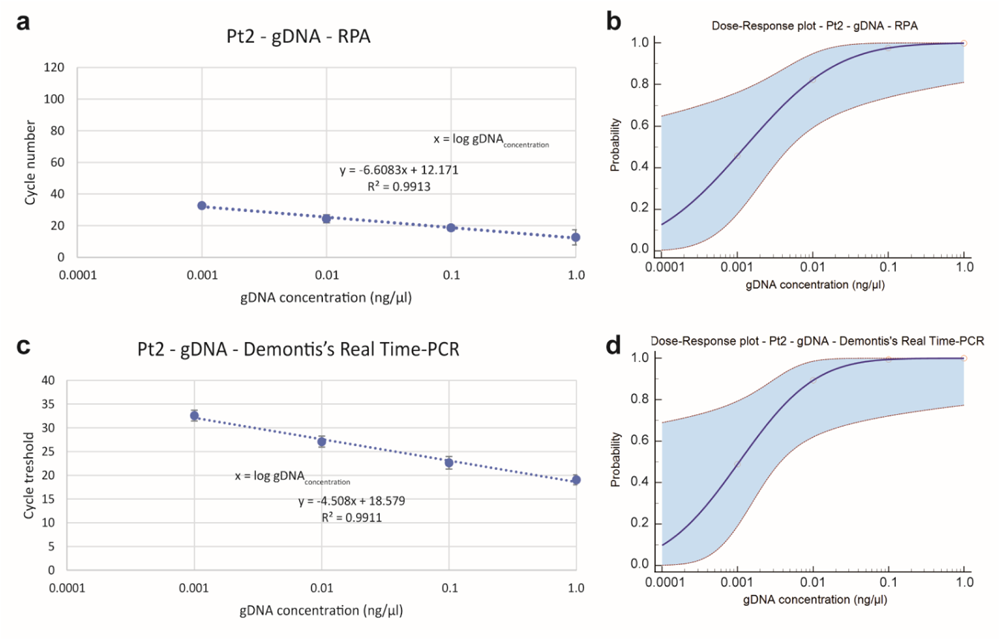
On the left, graphs showing the linear regression analysis of RPA (a) and Demontis’s Real Time-PCR (b) assays for the evaluation of their sensitivity toward independent serial dilutions of purified gDNA of the *P. tracheiphilus* isolate Pt2; bars indicate standard deviation (SD); blue dashed lines are the linear regression curves (linear equations and R^2^ values are reported). On the right, Dose-Response probit regression analyses showing the probability of detection with RPA (c) and Demontis’s Real Time-PCR (d) assays.

The probit regression “Dose-Response” analysis at 95% of probability and confidence interval evidenced for RPA a LoD of about 0.048 ng of gDNA (confidence interval ‒ CI ‒ range: 1.02 × 10^-2^ ‒ 8.049 × 10^3^ ng of gDNA), while for Demontis’s Real Time-PCR a LoD of about 0.02 ng of gDNA (CI range: 5.27 × 10^-3^ ‒ 1.602 × 10^7^ ng of gDNA) (Figure 3, b and d).

In ‘procedure ii’ (Figure 2b) the gDNA of 10^9^ conidia of the isolate Pt2 was extracted in accordance with the rapid DNA extraction protocol of (Edwards *et al*., 1991). Then, the sensitivity was evaluated on 10-folds dilutions calibrated at a gDNA concentration in the range 1.0 to 0.0001 ng/μL (2.85 × 10^3^ ‒ 0.285 genome copies/µl). Again, both RPA and Demontis’s Real Time-PCR gave amplifications up to 0.001 ng of gDNA (*i.e.* 1.0 pg of gDNA, corresponding to ∼ 29 genome copies/reaction) (Figure 4, a and c).

**Figure 4.**
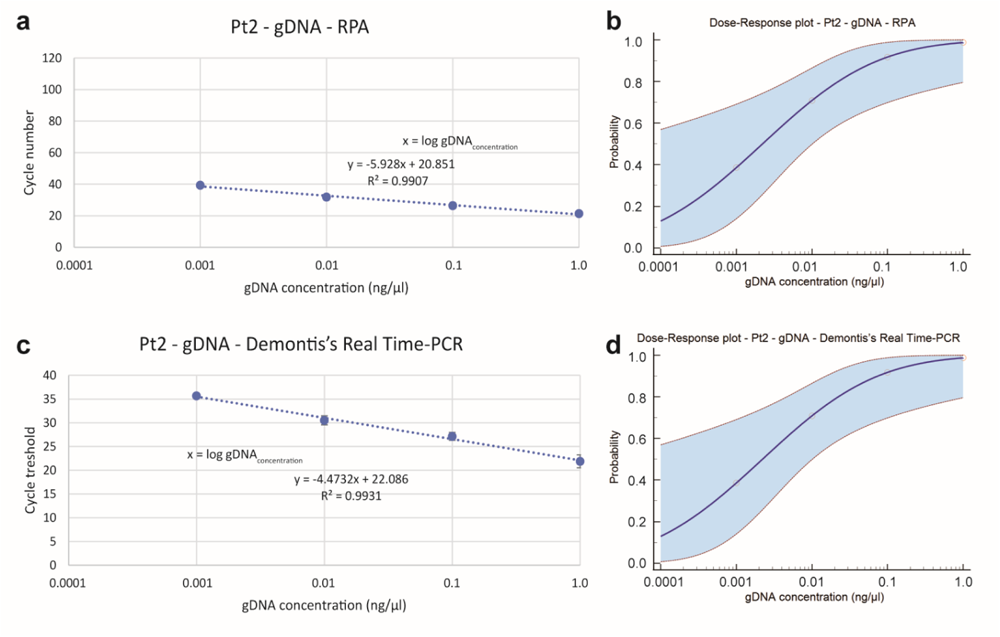
On the left, graphs showing the linear regression analysis of RPA (a) and Demontis’s Real Time-PCR (b) assays for the evaluation of their sensitivity toward independent serial dilutions of non-purified gDNA of the *P. tracheiphilus* isolate Pt2; bars indicate SD; blue dashed lines are the linear regression curves (linear equations and R^2^ values are reported). On the right, Dose-Response probit regression analyses showing the probability of detection with RPA (c) and Demontis’s Real Time-PCR (d) assays.

With this procedure the LoD was about 0.20 ng of gDNA (CI range: 3.26 × 10^-2^ ‒ 2.42 × 10^3^ ng of gDNA) for both RPA and Demontis’s Real Time-PCR (Figure 4, b and d).

In ‘procedure iii’ (Figure 2c) 10-folds suspensions of conidia of Pt2, from 10^8^ to 10 conidia/mL, in GEB lysis buffer were subjected to rapid DNA extraction performed in accordance (Edwards *et al*., 1991). Results of comparison of the two detection methods revealed similar sensitivity for both RPA and Demontis’s Real Time-PCR, with amplifications, for both techniques, in samples that were at the concentration from 10^8^ to 10^4^ conidia/ml (Figure 5, a and c).

**Figure 5.**
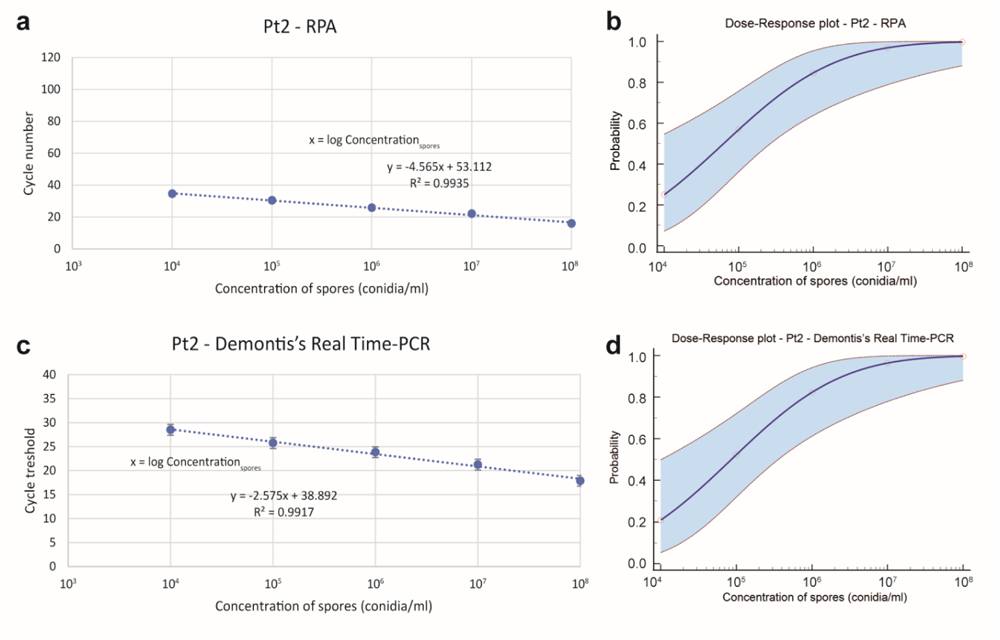
On the left, graphs showing the linear regression analysis of RPA (a) and Demontis’s Real Time-PCR (b) assays for the evaluation of their sensitivity toward non-purified gDNA of the *P. tracheiphilus* isolate Pt2 obtained from 10-folds conidia suspensions (from 10^8^ to 10 conidia/mL) in GEB lysis buffer; bars indicate SD; blue dashed lines are the linear regression curves (linear equations and R^2^ values are reported). On the right, Dose-Response probit regression analyses showing the probability of detection with RPA (c) and Demontis’s Real Time-PCR (d) assays.

For RPA the LoD was about 5.49 × 10^6^ conidia/ml (CI range: 9.22 × 10^5^ – 1.99 × 10^9^ conidia/ml), while for Demontis’s Real Time-PCR the LoD was about 6.7 × 10^6^ conidia/ml (CI range: 1.16 × 10^6^ ‒ 1.59 × 10^9^ conidia/ml) (Figure 5, b and d).

‘Procedure iv’ (Figure 2d) aimed at verifying the influence of plant matrix (twig cuttings and petioles) on the RPA sensitivity. To this aim, 10-folds suspensions of conidia of Pt2 (from 10^8^ to 10 conidia/mL) were established in a crude plant extract of either twig cutting or petioles collected from mal secco-free lemon plants. Results from samples suspended in plant crude extract from twig cuttings highlighted for both RPA and Demontis’s Real Time-PCR amplifications in the range 10^8^ - 10^4^ conidia/ml (Figure 6, a and c).

**Figure 6.**
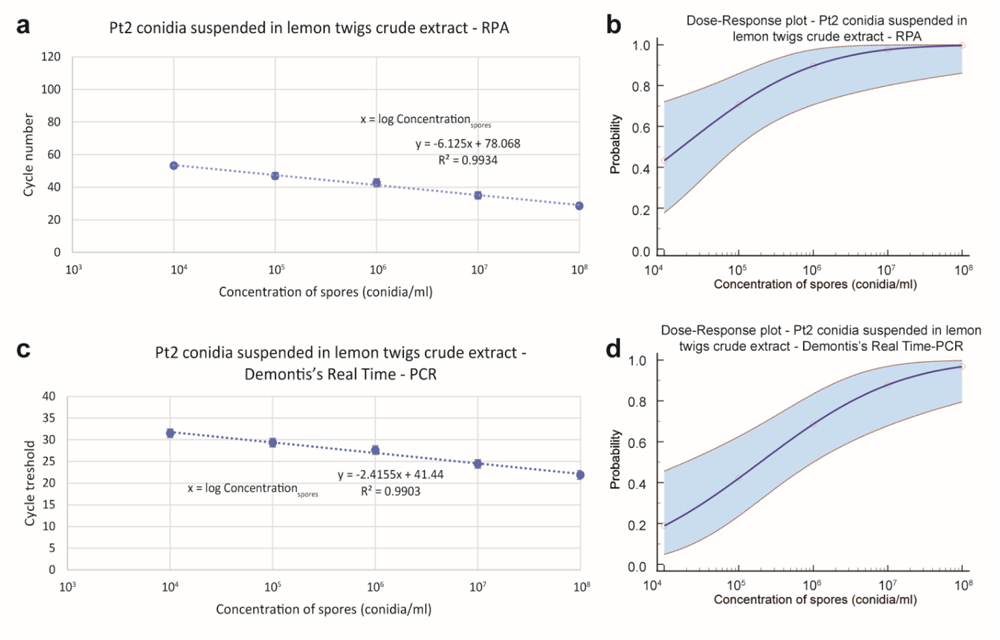
On the left, graphs showing the linear regression analysis of RPA (a) and Demontis’s Real Time-PCR (b) assays for the evaluation of their sensitivity toward non-purified gDNA of the *P. tracheiphilus* isolate Pt2 obtained from 10-folds conidia suspensions (from 10^8^ to 10 conidia/mL) in lemon twigs crude extract; bars indicate SD; blue dashed lines are the linear regression curves (linear equations and R^2^ values are reported). On the right, Dose-Response probit regression analyses showing the probability of detection with RPA (c) and Demontis’s Real Time-PCR (d) assays.

In this experimental condition RPA LoD was about 3.46 × 10^6^ conidia/ml (CI range: 4.72 × 10^5^ ‒ 4.03 × 10^10^ conidia/ml), while the Demontis’s Real Time-PCR LoD was about 4.97 × 10^7^ conidia/ml (CI range: 3.21 × 10^6^ ‒ 3.68 × 10^10^ conidia/ml) (Figure 6, b and d). With reference to samples obtained by the suspension of conidia in plant crude extract from petioles, both RPA and Demontis’s Real Time-PCR had positive amplifications in the range 10^8^ - 10^4^ conidia/ml (Figure 7, a and c); in this experiment RPA LoD was about 4.30 × 10^6^ conidia/ml (CI range: 1.07 × 10^6^ ‒ 9.48 × 10^9^ conidia/ml), while the Demontis’s Real Time-PCR LoD reached 2.29 × 10^7^ conidia/ml (CI range: 4.49 × 10^6^ - 1.01 × 10^10^ conidia/ml) (Figure 7, b and d).

**Figure 7.**
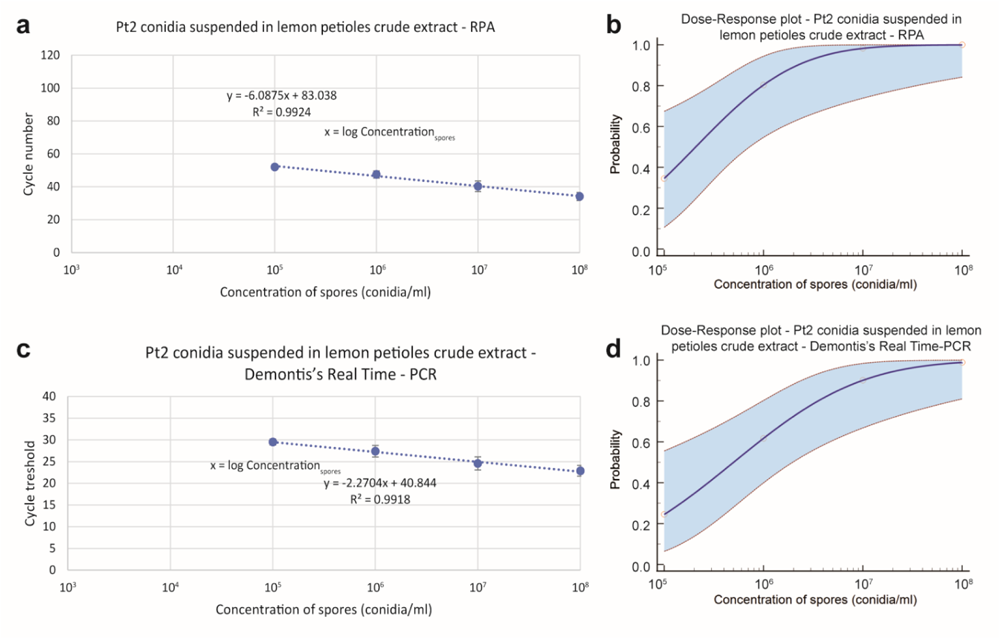
On the left, graphs showing the linear regression analysis of RPA (a) and Demontis’s Real Time-PCR (b) assays for the evaluation of their sensitivity toward non-purified gDNA of the *P. tracheiphilus* isolate Pt2 obtained from 10-folds conidia suspensions (from 10^8^ to 10 conidia/mL) in lemon petioles crude extract; bars indicate SD; blue dashed lines are the linear regression curves (linear equations and R^2^ values are reported). On the right, Dose-Response probit regression analyses showing the probability of detection with RPA (c) and Demontis’s Real Time-PCR (d) assays.

### 2.3. Efficiency of developed RPA assay in detecting P. tracheiphilus DNA in symptomatic and asymptomatic plant material

In order to evaluate the practical application of the developed RPA assay, eight samples of *P. tracheiphilus*- susceptible plant material (twigs and petioles) from mal secco-symptomatic and -non symptomatic lemon plants were collected from local commercial lemon orchards and tested for the presence of the pathogen using classical isolation, Demontis’s Real Time-PCR and RPA amplification (Table 3). Successful outcomes by the three detection techniques were reported from all twig samples collected from symptomatic plants (ID01, ID04, ID07; Table 3), with Demontis’s Real Time-PCR cycle threshold in the range 27.11-28.81 (Table 3). With reference to twig samples collected from non-symptomatic plants (ID03 and ID08), ID03 resulted positive just to the molecular detection methods, whereas ID08 resulted negative to *P. tracheiphilus*-detection with all the techniques. Among samples of petioles collected from symptomatic plants, ID02 and ID06 (Table 3), ID02 was *P. tracheiphilus*-positive just by RPA, while ID06 gave positivity to the presence of *P. tracheiphilus* by all the employed detection techniques (Demontis’s Real Time-PCR cycle threshold, 25.51). Finally, no detection by any technique was reported from the petiole sample ID05 collected from a non-symptomatic host (Table 3).

**Table 3.** Results from detection tests carried out on plant matrices (twigs/petioles) collected from mal secco-symptomatic and -non symptomatic lemon plants.

| Sample ID | Source of sample | signs/symptoms on the plant (P<sup>1</sup>/A<sup>2</sup>) | Sample (lemon twig/petiole) | Location | *P. tracheiphilus* isolation outcome (+<sup>3</sup>/-<sup>4</sup>) | Demontis's Real Time-PCR (Cycle threshold) | RPA outcome (+<sup>3</sup>/-<sup>4</sup>) |
| --- | --- | --- | --- | --- | --- | --- | --- |
| ID01 | *Citrus × limon* | P | Twig | Syracuse (Sicily) | + | 28.81 | + |
| ID02 | - | P | Petiole | Syracuse (Sicily) | - | nd<sup>5</sup> | + |
| ID03 | - | A | Twig | Avola (Syracuse, Sicily) | - | 31.49 | + |
| ID04 | - | P | Twig | Avola (Syracuse, Sicily) | + | 26.46 | + |
| ID05 | - | A | Petiole | Mineo (Catania, Sicily) | - | nd | - |
| ID06 | - | P | Petiole | Mineo (Catania, Sicily) | + | 25.51 | + |
| ID07 | - | P | Twig | Mineo (Catania, Sicily) | + | 27.11 | + |
| ID08 | - | A | Twig | Mineo (Catania, Sicily) | - | nd | - |

<sup>1</sup> P = presence; <sup>2</sup> A = absence; <sup>3</sup> + = positive; <sup>4</sup> - = negative; <sup>5</sup> nd = no-detection by Real Time-PCR.

## 3. Discussion

Phytosanitary emergencies require tools capable of reducing the time necessary for decision-making processes to rapidly develop the most effective management strategies (Cacciola and Gullino, 2019). This is particularly relevant for quarantine plant pathogens, such as *P. tracheiphilus*, whose management often involves the adoption of strict measures that limit the trade of plant products and propagation material (Migheli *et al*., 2009). For this reason, nowadays, these emergencies must rely on technologically advanced means, such as molecular techniques able to satisfy the major requirements of the detection, i.e. specificity, sensitivity and rapidity (Aslam *et al*., 2017; Boonham *et al*., 2016; Donoso and Valenzuela, 2018; Hariharan and Prasannath, 2021; Marcolungo *et al*., 2022; Patel *et al*., 2022).

The most reliable diagnostic protocols so far developed for the early detection of *P. tracheiphilus* in possibly infected plant material were officially issued by the European Plant Protection Organization (EPPO) and were all based on different molecular approaches. The first molecular standard issued by EPPO for the diagnosis of *P. tracheiphilus* (EPPO, 2005) was based on studies carried out by Rollo *et al*. (1987, 1990), and employed a dot blot approach in combination with the polymerase chain reaction (PCR). Advancements in DNA sequencing and bioinformatic tools have fostered the search for highly specific protocols for the detection of plant pathogens (Venbrux *et al*., 2023). The study by Balmas *et al*. (2005) achieved a very high diagnostic specificity resulting in a very reliable detection of *P. tracheiphilus* causing mal secco. Specifically, in Balmas *et al*. (2005), the phylogenetic characterization of a population of Italian isolates of *P. tracheiphilus* provided several ITS1-5.8S-ITS2 sequences. The alignment of such sequences highlighted a 544 bp highly specific consensus sequence that provided the basis for the design of the primer pair Pt-FOR2/Pt-REV2. The high *P. tracheiphilus*-specificity of these primers made them an obvious choice in the updated version of the 2015 of the EPPO diagnostic protocol (EPPO, 2015). Other primers targeting the ITS region for the detection of *P. tracheiphilus* in infected tissues of the host plant by conventional PCR were subsequently developed by other authors (Ezra *et al*., 2007; Kalai *et al*., 2010). The performance of these primers was comparable to that of the primers designed by (Balmas *et al*., 2005). Despite its high versatility, conventional PCR is often not enough sensitive to guarantee plant material as being free from the pathogen; additionally, it is not quantitative (Demontis *et al*., 2008). More recently, the development of real time PCR, also known as qPCR, has resulted in protocols with improved sensitivity for the detection of *P. tracheiphilus* (Demontis *et al*., 2008; Licciardello *et al*., 2006). The TaqMan® / SYBR® Green I real-time PCR protocol of Demontis *et al*. (2008) was included as quantitative method in the PM 7/048 (3) *Plenodomus tracheiphilus* EPPO diagnostic protocol (EPPO, 2015), due to its reliability and high sensitivity (detecting as little as 15 pg of the target pathogen’s DNA).

Although the above protocols meet the requirements for specificity and sensitivity, their major drawback remains the time necessary to complete the assay. In fact, in addition to the technical time needed to complete the PCR run (which usually ranges between 1,30-1,45 hours) and to analyze results, DNA samples that will undergo amplification by *Taq* polymerase require high purity (Demeke and Adams, 1992; Demeke and Jenkins, 2010; Henson and French, 1993; John, 1992; Schrader *et al*., 2012; Sipahioglu *et al*., 2006; Su and Gibor, 1988; Wan and Wilkins, 1994; Wei *et al*., 2008). The need for pure DNA extracts extend the timing of the detection by PCR techniques to about one working day (Demontis *et al*., 2008). Isothermal amplification technologies, such as loop-mediated amplification (LAMP), helicase-dependent amplification (HDA), nucleic acid sequence-based amplification (NASBA), rolling circle amplification (RCA) and recombinase polymerase amplification (RPA) have shortened the time necessary to achieve a diagnostic outcome (Ivanov *et al*., 2021). These methods differ from each other in terms of amplification temperatures, sensitivities, reaction times, and other advantages and drawbacks (Gill and Ghaemi, 2008; Ivanov *et al*., 2021). More recently, diagnostic assays based on RPA have become widely used for the molecular-based diagnosis of diseases (Tan *et al*., 2022). The outbreak of SARS-CoV-2 has further promoted the application of RPA in nucleic acid detection (Bai *et al*., 2022). Previous studies on RPA detection have shown several advantages of this technique, including its user-friendliness (in terms of simplicity of operations and low equipment requirements), high sensitivity, and great specificity (Tan *et al*., 2022). Additionally, RPA best matches the requirement of ‘rapidity’. In this respect, with a reaction time of no more than 20 minutes, RPA is not only the fastest isothermal amplification technique, but it is also the most reliable technique in terms of tolerance to inhibitors of the molecular reaction (Boluk *et al*., 2020; Chandu *et al*., 2016; Kapoor *et al*., 2017). Inhibition of the molecular reaction is one of the major concerns in PCR technologies (Sidstedt *et al*., 2020). A PCR inhibitor is defined as any substance that is capable of interfering with one or more of the molecular reactions involved in PCR (denaturation, primer annealing, binding of polymerase to primer-DNA complex, primer extension, and fluorescence - in real-time technologies) and that consequently determines an inhibition of the amplification (Hedman *et al*., 2013). PCR inhibitors comprise a wide array of substances, including inorganic ions, organic salts, organic acids, some dyes and other molecules (Hedman *et al*., 2013). The wide tolerance of RPA to common PCR inhibitors is a highly relevant strength of this technique, because DNA samples obtained through rapid extractions, although not purified from inhibitors, can be used in RPA amplifications with a high chance of success (Ivanov *et al*., 2021). This is the first study describing the development of a molecular diagnostic assay for the mal secco of lemon caused by *P. tracheiphilus* that is based on isothermal amplification of nucleic acids. These tests confirmed the exclusive amplification of the selected barcode of the target organism and led to the design of the final element of this detection assay, the RPA probe (RPA_Ptrach_Probe; Table 1). Then, RPA runs made it possible to validate the whole *P. tracheiphilus-*specificity and -inclusivity of the developed RPA assay. In this study, four different sensitivity tests were carried out to highlight the strengths and limitations of the developed RPA assay compared to the SYBR® Green I Real Time-PCR test by Demontis *et al*. (2008). Results highlighted that the RPA assay was as sensitive as Demonti’s Real Time-PCR under the same conditions. Both techniques reported a detection threshold of 1.0 pg of gDNA per reaction (corresponding to about 29-genome copies/reaction) and a 95% of probability LoD of order of magnitude 10 pg gDNA/reaction.

In order to further investigate the performances of the RPA assay and to highlight its potential for the diagnosis of the disease in the field, specific validation trials were performed. Under our experimental conditions, RPA was ten times more sensitive than Demonti’s Real Time-PCR when testing conidial DNA extracts. A one order of magnitude higher sensitivity of the RPA compared to Demonti’s assay was also confirmed when crude plant extracts, and supposedly inhibitors, were added to the substrate tested. This result is consistent with previous studies that have demonstrated the high reliability of RPA in amplification runs in the presence of plant macerates (Boluk *et al*., 2020; Chandu *et al*., 2016; Kapoor *et al*., 2017), as well as with other studies that highlighted a significant inhibition of PCR in the presence of plant metabolites such as pectin, polyphenols, polysaccharides, and xylan (Demeke and Adams, 1992; Henson and French, 1993; John, 1992; Monteiro *et al*., 1997; Schrader *et al*., 2012; Sipahioglu *et al*., 2006; Su and Gibor, 1988; Wan and Wilkins, 1994; Wei *et al*., 2008).

The effectiveness of the RPA assay was finally evaluated for detecting mal secco infections *in planta*. The sensitivity of the assay was compared to Demontis’s Real Time-PCR and classical pathogen isolation. The two molecular detection approaches were superior to the isolation approach and comparable to each other. In conclusion, the RPA assay’s specificity and inclusivity were validated through various tests, and it was found that RPA was more sensitive than the Real Time-PCR test. However, it is important to note that the accuracy and reproducibility of any diagnostic tool are critical for its effective use. Therefore, results obtained here deserve to be validated through ring tests, which involve multiple laboratories performing the same assay on the same samples. The RPA assay has several advantages over traditional detection methods, including its speed, ease of use, and potential for field applications. Based on the promising result presented in this study, further research is needed to optimize and validate this technique for use in various conditions and to make it more accessible to farmers and researchers.

## 4. Experimental procedures

### 4.1. Fungal isolates

Fungal and oomycete isolates employed in this study belonged to the collection of the Laboratory of Molecular Plant Pathology of the Department of Agriculture, Food and Environment of the University of Catania (Catania, Italy). Test isolates included a wide collection of virulent strains of *P. tracheiphilus*, *Pleosporales* isolates from species taxonomically close to *P. tracheiphilus*, and several specimens of the major fungal and oomycete species commonly associated to citrus (Table 2); these latter comprised: *Phytophthora* spp. (*P. nicotianae* and *P. citrophthora*), *Alternaria* spp. (*A. alternata* and *A. arborescens*), *Colletotrichum* spp. (*C. acutatum*, *C. gloeosporioides* and *C. karsti*), *Penicillium* spp. (*P. digitatum* and *P. italicum*), *Phyllosticta citricarpa*. Most of them had been characterized at species level in previous studies (El boumlasy *et al*., 2021; Demontis *et al*., 2008; Riolo *et al*., 2021; Rovetto *et al*., 2023). Specimens of fungal organisms obtained in this study (Table 2) were molecularly characterized by PCR amplification, analysis and sequencing of specific barcode regions. In detail, isolates of *P. tracheiphilus* were characterized on the basis of sequences of the internal transcribed spacer (ITS) region amplified by using primer pairs ITS1/ITS4 (White *et al*., 1990). For *Penicillium* isolates, the barcodes were the ITS region (amplified as above) and the β-tubulin gene (*tub2*), this latter amplified by using primer pairs Bt2a/Bt2b (Glass and Donaldson, 1995). For *Phyllosticta citricarpa,* the used barcode was part of the translation elongation factor 1-α gene (*tef1*), amplified by using primer pairs EF1-728F/EF2 (Carbone and Kohn, 1999; O’Donnell *et al*., 1998). PCR reactions were performed using the *Taq* DNA polymerase recombinant (Invitrogen™, Carlsbad, 254 CA, USA) following the manufacturer’s instructions.

All fungal and oomycete isolates were maintained at the temperature of 25°C on Potato Dextrose Agar (PDA).

### 4.2. Selection of Plenodomus tracheiphilus target region and development of RPA primers and probe

A 142 bp throughout barcode belonging to the Internal Transcriber Spacer (ITS) regions of the ribosomal DNA (rDNA) of *P. tracheiphilus* (Figure 1) was selected by alignment in MEGAX (MEGA – Molecular Evolutionary Genetics Analysis) of several NCBI deposited ITS sequences (Genbank accession numbers: AY531665, AY531666, AY531667, AY531668, AY531669, AY531670, AY531671, AY531672, AY531673, AY531674, AY531675, AY531676, AY531677, AY531678, AY531679, AY531680, AY531681, AY531682, AY531689) belonging to officially identified isolates of *P. tracheiphilus* (Balmas *et al*., 2005). Barcode specificity was confirmed by BLAST searches on NCBI nucleotide database (Home - Nucleotide - NCBI). Primers for the RPA amplification of the 142 bp selected barcode were designed by using the Primer BLAST NCBI tool (Primer designing tool) (Table 1). *Plenodomus tracheiphilus-*specificity of designed primers was preliminarily confirmed by both *in silico* amplification (Primer BLAST NCBI tool was used) and conventional PCR; this latter was performed on 1 ng of genomic DNA (gDNA) of each organism listen in Table 2 by using the *Taq* DNA polymerase recombinant (Invitrogen™, Carlsbad, 254 CA, USA) following the manufacturer’s instructions. The exclusive PCR amplification of gDNA from isolates of *P. tracheiphilus* (Table 2) was highlighted by electrophoresis in TAE 1X Agarose gel; then, *P. tracheiphilus* PCR products were sequenced by an external service (Macrogen, Seoul, South Korea) and those sequences, which totally matched the 142 bp of the selected barcode, were aligned by MEGAX (MEGA - Molecular Evolutionary Genetics Analysis).

The RPA probe was designed on a 50 bps fragment within the selected barcode (Figure 1); probe was labeled by a tetrahydrofuran residue (THF), flanked by a dT-fluorophore (FAM) and a corresponding dT-quencher (dt-Q) group; additionally, the probe was blocked at the end 3′ by a C3-Spacer (C3-S) modification group (Table 1).

### 4.3. RPA amplifications

Each RPA reaction performed in this study was carried out in a 0.25 mL tube (AmplifyRP Discovery kits Agdia Inc., Elkhart, IN, United States) containing RPA mastermix pelleted reagents, a rehydration solution (rehydration buffer - 68.6% v/v, forward and reverse primers - 0.40 μM, XRT probe - 0.11 μM, and nuclease free water till the volume of 21.5 ul), MgOAc (1.25 μL) and gDNA (1.0 μL). Amplifications were run in the AmpliFire® Isothermal Fluorometer (Agdia Inc., Elkhart, IN, United States); conditions consisted of a 20 minutes heating at the constant temperature of 39°C. The amplification curve files were used for constructing linear regression curves and statistical analyses. Each RPA run included negative (nuclease free water) and positive (1 ng of gDNA of the isolate Pt2) controls that were used for samples comparisons. Each reaction was carried out in triplicate.

### 4.4. Evaluation of P. tracheiphilus-specificity and -inclusivity of RPA assay

Specificity of RPA assay was checked against its plant host (*Citrus limon*) and a panel of selected fungal and oomycete isolates (Table 2). Samples were represented by stem fragments from *Citrus limon* cv. Femminello Siracusano and, for fungal and oomycete pathogens, by fresh mycelium collected from 7-days-old cultures grown on PDA at 25°C. For both plant tissues and fungal and oomycete microorganisms the genomic DNA (gDNA) was extracted by using the PowerPlant^®^ Pro DNA Isolation Kit (the manufacturer’s instructions were followed). For each gDNA sample, the concentration was checked by using the Qubit fluorimeter (Invitrogen) and adjusted at 1 ng/μL. RPA reactions were performed in triplicate and carried out as described in 2.3. Inclusivity of RPA assay was checked toward DNA isolated from 29 specimens of *P. tracheiphilus* (Table 2); DNA samples preparation and RPA runs were as above.

### 4.5. Evaluation of sensitivity of the RPA assay vs. Demontis’s Real Time-PCR assay

Tests for the evaluation of the sensitivity were conducted on gDNA of the isolate *P. tracheiphilus* Pt2. Sensitivity of the developed RPA assay in detecting gDNA of *P. tracheiphilus* was tested in comparison with the sensitivity of the SYBR^®^ Green I Real Time-PCR test of Demontis *et al*. (2008). Each Real Time-PCR reaction consisted of 10 μl of PowerUp^TM^ SYBR^TM^ Green Master Mix (2×) (Applied Biosystems^TM^, Foster City, CA, United States), 0.5 μM of each primer (Phomafor: 5′-GCTGCGTCTGTCTCTTCTGA-3′; Phomarev: 5′-GTGTCCTACAGGCAGGCAA-3′), 1 ul of DNA template and nuclease free water till the final volume of 20 uL. Each run included negative (nuclease free water) and positive (1 ng of gDNA of the isolate Pt2) controls that were used for samples comparisons. All Real Time-PCR reactions were carried out in triplicate.

The sensitivity of RPA assay was evaluated toward different kinds of samples from which DNA was isolated through different kinds of procedures (Figure 2).

#### Procedure i

10^9^ conidia of the isolate Pt2 were subjected to gDNA extraction by using the PowerPlant® Pro DNA Isolation Kit (Qiagen, Hilden, Germany) (the manufacturer’s instructions were followed); the concentration of the obtained DNA sample was checked by Qubit fluorometer (Invitrogen^TM^, Waltham, MA, United States); then, 10-folds dilutions (from 1.0 to 0.0001 ng/μL) in 1X TE buffer (10 mM Tris–HCl and 1 mM EDTA; pH 8.0) were realized. All the obtained samples were stored at -20°C until molecular analyses (Figure 2a).

#### Procedure ii

10^9^ conidia of the isolate Pt2 were suspended in 1 mL of GEB lysis buffer (Agdia Inc., Elkhart, IN, United States) and the mixture was grinded by a konte pestle until a satisfactory liquefaction was observed; then, the extraction of DNA was carried out in accordance with the rapid DNA extraction protocol of Edwards *et al*. (1991) with slights modifications: after liquefaction, 1 mL of isopropanol was added. The sample was vortexed for 5 seconds and centrifuged at 13,000 rpm for 5 minutes. After centrifugation, the supernatant was discharged and the obtained pellet was air-dried for 5 minutes. Then, the pellet was dissolved in 100 μL of 1X TE buffer (10 mM Tris–HCl and 1 mM EDTA; pH 8.0) and the obtained solution was centrifuged at 13,000 rpm for 1 minute. After centrifugation, the supernatant was transferred into a new 1.5 ml tube. The concentration of the obtained DNA sample was checked by Qubit fluorometer (Invitrogen^TM^, Waltham, MA, United States); then, 10-folds dilutions (from 1.0 to 0.0001 ng/μL) in 1X TE buffer (10 mM Tris–HCl and 1 mM EDTA; pH 8.0) were realized. All the obtained samples were stored at − 20°C until molecular analyses (Figure 2b).

#### Procedure iii

2.0 x 10^8^ conidia of the isolate Pt2 were suspended in 2 mL of GEB lysis buffer (Agdia Inc., Elkhart, IN, United States) and 10-folds dilutions (from 10^8^ to 10 conidia/mL) in the same buffer, to a final volume of 1 mL, were realized. Each 1 mL sample of conidia in GEB lysis buffer (concentrations from 10^8^ to 10 conidia/mL) was grinded by a konte pestle until a satisfactory liquefaction was observed; then, for each sample, the extraction of the DNA proceeded in accordance with the rapid DNA extraction protocol of Edwards *et al*. (1991) with slights modifications, as reported in Procedure ii (Figure 2c).

#### Procedure iv

in order to verify the influence of plant matrix (twig cuttings and petioles) on the RPA sensitivity, a specific rapid gDNA extraction procedure was accomplished (Figure 2d).

#### STEP 1 – Preparation of a crude plant extract - plant tissue amended GEB lysis buffer (GEB – lemon)

mal secco-free twig cuttings (or petioles) were powdered by using a cheese grater; then, the obtained powder was added to GEB lysis buffer (Agdia Inc., Elkhart, IN, United States) (proportion 1 g of powder:10 ml of lysis buffer); the mixture was vortexed for 5 seconds and followed a centrifugation round consisting of 13,000 rpm for 5 minutes; the obtained supernatant was transferred to new 1.5 ml tubes and employed as lysis buffer in the following described gDNA extraction.

#### STEP 2 – gDNA extraction

2.0 x 10^8^ conidia of the isolate Pt2 were suspended in 2 mL of GEB – lemon (obtained from twig cuttings or petioles) and 10-folds dilutions (from 10^8^ to 10 conidia/mL) in GEB – lemon, to a final volume of 1 mL, were realized. Each 1 mL sample of conidia in GEB – lemon (concentrations from 10^8^ to 10 conidia/mL) was grinded by a konte pestle until a satisfactory liquefaction was observed; then, for each sample, the extraction of the DNA proceeded in accordance with the rapid DNA extraction protocol of Edwards *et al*. (1991) with slights modifications, as reported in Procedure ii.

Linear regression curves of RPA and Real Time-PCR were calculated by using Excel software. The Limit of Detection (LoD) of both RPA and Real Time PCR assays at 95% of probability and confidence interval was calculated by Probit regression “Dose-Response” analysis performed with eight replicates by using MedCalc Software (Ostend, Flanders, 8400, Belgium). In agreement with the method followed by Cesbron *et al*. (2023), in this study the genome copy number was calculated by using the estimated genome size of the reference isolate *P. tracheiphilus* strain IPT5 (34,242,632 bp; GenBank accession number: GCA_010093695.1), knowing that the mean weight of one nucleotide pair is 1.023 × 10^-9^ pg (Dolezel *et al*., 2003).

### 4.6. Evaluation of the efficiency of the RPA assay in detecting Plenodomus tracheiphilus DNA in symptomatic and asymptomatic plant material

In order to evaluate the practical application of the RPA assay in comparison with Demontis’s Real Time-PCR, tests were carried out on Mal secco-symptomatic and -asymptomatic plant material (lemon twigs and petioles); samples were collected in October 2022 from commercial lemon orchards of the area of Catania and Siracusa (Italy) (Table 3). For each sample, total DNA was extracted as it follows: 0.1 g of plant material (lemon twigs or petioles) were powdered by using a cheese grater, added to 1 mL of GEB lysis buffer (Agdia Inc., Elkhart, IN, United States) and grinded by a konte pestle until a satisfactory liquefaction was observed; then, 1 mL of isopropanol was added, the sample was vortexed for 5 seconds and centrifuged at 13,000 rpm for 5 minutes; after centrifugation, the supernatant was discharged and the obtained pellet was air-dried for 5 minutes; then, the pellet was dissolved in 100 μL of 1X TE buffer (10 mM Tris–HCl and 1 mM EDTA; pH 8.0) and the obtained solution was centrifuged at 13,000 rpm for 1 minute; after centrifugation, the supernatant was transferred into a new 1.5 ml tube.

Positivity to Mal secco was additionally verified by isolations and molecular detection of *P. tracheiphilus* isolates; this latter was performed by conventional PCR associated to electrophoresis and UV observation of PCR products, in accordance with the PM 7/048 *P. tracheiphilus* EPPO diagnostic protocol (Balmas *et al*., 2005; EPPO, 2015). For isolations, plant material (girdled tissues or petioles) was cut into small pieces (3.0-4.0 mm in diameter) and subjected to a surface sterilization consisting on immersion in 1% NaClO for 1 min., rinsing in sterilized distilled water (sdw) (1 min.), immersion in 70% ethanol (1 min.) and final rinsing in sterilized distilled water (1 min.). Sterilized pieces were blotted dry with adsorbent paper, plated onto Potato Dextrose Agar (PDA) amended with streptomycin sulphate (250 mg/L) and incubated at 22-23°C for 48 hours. After incubation, outgrowing hyphas were transferred on PDA plates and let to growth at 22-23°C for 10-12 days. Then, the final isolates were obtained by monoconidial culture.

## Accession numbers

Barcode sequences of isolates obtained in this study were submitted to the GenBank database under the following accession numbers: *Plenodomus tracheiphilus* isolate Pt22 (OR656741), Pt3 (OR656742), Pt6-Lim-Sic (OR656743), Pt8 (OR656744), Pt9-Lim-Sic (OR656745), Pt10 (OR656746); *Phyllosticta citricarpa* isolate 1G (OR665395); *Penicillium italicum* isolate T4N0 (OR652459, OR665396).

## Author Contributions

SOC, FLS, MA, MG and SM conceptualized the study, analyzed the results, and reviewed and edited the draft. EIR and FLS did the investigation and formal analysis and performed the experiments. SOC, FLS and MA were responsible for funding acquisition and supervised the study. FLS and EIR wrote the original draft. SOC, MG and SM revised the draft of the manuscript. All authors contributed to the article and approved the submitted version.

## Acknowledgements

This research was funded by the University of Catania, Italy “Investigation of phytopathological problems of the main Sicilian productive contexts and eco-sustainable defense strategies (ME-DIT-ECO) PiaCeRi-PIAno di inCEntivi per la Ricerca di Ateneo 2020-22 linea 2” “5A722192155”, by the projects “Smart and innovative packaging, postharvest rot management, and shipping of organic citrus fruit (BiOrangePack)” under the Partnership for Research and Innovation in the Mediterranean Area (PRIMA)—H2020 (E69C20000130001), the “Italie–Tunisie Cooperation Program 2014–2020” project “PROMETEO «Un village transfrontalier pour protéger les cultures arboricoles méditerranéennes en partageant les connaissances»” cod. C-5-2.1-36, CUP 453E25F2100118000, and by the European Union (NextGeneration EU), through the MUR-PNRR project SAMOTHRACE (ECS00000022).

## Conflict of Interest Statement

MA was employed by the company Agdia EMEA.The remaining authors declare that the research was conducted in the absence of any commercial or financial relationships that could be construed as a potential conflict of interest.

## Notes

### Competing Interest Statement

Marcos Amato was employed by the company Agdia EMEA. The remaining authors declare that the research was conducted in the absence of any commercial or financial relationships that could be construed as a potential conflict of interest.

